# Liquid biopsy in gastrointestinal stromal tumors –simultaneous detection of primary and secondary mutations with an NGS based circulating tumor DNA assay

**DOI:** 10.1101/481622

## Introduction

Gastrointestinal Stromal tumors (GISTs) are the most common neoplasms of mesenchymal origin arising from the gastrointestinal tract. GISTs are mostly characterised by one predominant activating mutation in *KIT* (*KIT* proto-oncogene receptor tyrosine kinase) or *PDGFRA* (platelet derived growth factor receptor alpha) genes^1-3^. Under pharmacological pressure, one or more secondary mutations can develop in *KIT*, conferring resistance to tyrosine kinase inhibitor, imatinib. In imatinib-resistant GISTs, secondary mutations can occur in the ATP binding pocket, encoded by exons 13 and 14 or in the kinase activation loop, encoded predominantly by exon 17^4^. *In vitro* studies have demonstrated that sunitinib is preferentially more active against the mutations in the ATP binding pocket while regorafenib, has increased activity against mutations in the kinase activation loop^5^.

Radiological response evaluation is the standard of care at this point; there is currently no specific biomarker to monitor disease progression or to determine the development of resistance. Primary GIST mutation is routinely detected from FFPE tissue via sanger sequencing. While knowledge of secondary mutations could enable tailoring of post-imatinib treatment, re-biopsy of a progressive metastatic lesion is not routine. It is invasive and may not comprehensively reflect the entire evolving tumor biology upon disease progression. Detection of secondary mutations may be affected by intra- and inter-lesional tumor heterogeneity^6^.

Non-invasive profiling with ctDNA is an attractive means to monitor disease volume (using primary mutations) and survey (for secondary mutations).

ctDNA assays are being developed for disease-specific and pan-cancer applications In GISTs, allele specific PCR (polymerase chain reaction)^7^, BEAMing (Beads Emulsion Amplifications and Magnetics)^8^, droplet PCR^9^ and next generation sequencing (NGS) assays^10-13^ have been deployed for detection of mutations from plasma via circulating tumor DNA (ctDNA). Assays targeting single mutations are highly sensitive, but are not easily generalizable in GISTs, which is characterized by a wide diversity of different primary mutations across patients, often a large deletion with differing breakpoints. Furthermore, these assays cannot detect emergent secondary mutations which also differ between patients. Broad-based sequencing methods can cover more mutations, but the sequencing depth required for highly sensitive allele detection may be prohibitive for a large target. In this study, we sought to overcome these limitations by designing a set of multiplex amplicon-based sequencing assays that have specific coverage for a patient’s primary mutations as well as general coverage of potential secondary mutation hotspots.

We hypothesized that circulating tumor DNA can be used as a dynamic biomarker to monitor disease status. The primary objective of this study was to determine the performance of our NGS-assay in detecting primary circulating *KIT* or *PDGFRA* mutations in the plasma of patients and correlating it with disease course. The secondary objective was to detect emergence of secondary *KIT* mutations in these patients.

## Methods

### Amplicon assay development

We developed a set of nine multiplexed amplicon sequencing assays covering primary and secondary mutations in GISTs, with each assay being a set of six primer pairs. Each set included one primer pair targeting a single primary mutation hotspot in *KIT* or *PDGFRA*, and five common primer pairs targeting secondary mutation hotspots in *KIT*. Primer design, optimisation, PCR conditions, amplicon generation, sequencing and variant calling were done as previously described^14^ with minor changes. Namely, a) primer design using Primer3Web v4.0.0 (http://primer3.ut.ee/) allowed, for amplicons over known deletion hotspots to be longer than 150bp b) PCR annealing temperature was set to 58^°^C and c) read alignment parameters were tuned to favour mapping with long insertion/deletions instead of soft clipping (−L 500,500 -E 1,1 -A 2) and variant sites were not pre-specified. For each patient, based on the primary mutation in that patient, one of the multiplex-amplicon assays was selected for use. The rationale being that the primary mutation (from routine diagnostic tissue biopsy) provides information on the primary mutation (and initiation of treatment) and the circulating tumor DNA is used for disease monitoring and surveillance for secondary mutations.

### Lower limit of detection & linearity of amplicon assay

Twelve GIST samples with mutations spanning our assays were selected. Using tumor DNA, the known primary mutation was confirmed using our assay. Subsequently, each DNA sample was separately mixed with HapMap NA18532 reference DNA in fixed proportions by mass to create a dilution series (1:1, 1:5, 1:10, 1:50, 1:100) with known mutation allele frequencies.

### Plasma processing and DNA extraction

Plasma and DNA extraction were performed as previously described^14^. In brief, blood specimens were processed within 2 hours of collection and plasma was extracted and stored at −80^°^C. Circulating DNA was subsequently extracted from plasma using the QiaAmp Circulating Nucleic Acids Kit (Qiagen). Based on the patient’s primary mutation as determined from clinical sequencing, the appropriate multiplex assay was selected for use on the plasma DNA.

### Assay performance

We designed and optimised a set of 9 multiplex-amplicon sequencing assays. Together, the assays cover 94% of known variants from the COSMIC variant database in the *KIT* and *PDGFRA* exons targeted (Table 1). The assays were first tested on 16 GIST tumor FFPE samples (formalin fixed, paraffin embedded tissue) with 100% concordance with the expected mutation from clinical sequencing. Separate serial dilution experiments demonstrated the ability of the assay system to detect mutations down to a variant allele frequency (VAF) of 0.2% with a strong positive linear correlation between the expected and observed VAF (R^2^=0.85, Fig 1)^15^.

**Table 1:** Overall mutational coverage of our GIST assay based on the COSMIC variant database (20180306)

| Gene | Exon | Covered | Total | Percentage | Primer Set |
| --- | --- | --- | --- | --- | --- |
| KIT | 9 | 542 | 678 | 79.94% | G01 |
| KIT | 11 | 3842 | 3846 | 99.90% | G03, G04, G10, G11 |
| KIT | 13 | 321 | 373 | 86.06% | All primer sets |
| KIT | 14 | 47 | 76 | 61.84% | All primer sets |
| KIT | 15 | 14 | 29 | 48.28% | All primer sets |
| KIT | 17 | 2522 | 2598 | 97.07% | All primer sets |
| KIT | 18 | 25 | 109 | 22.94% | All primer sets |
| PDGFRA | 12 | 170 | 216 | 78.70% | G06 |
| PDGFRA | 14 | 60 | 74 | 81.08% | G07 |
| PDGFRA | 18 | 896 | 980 | 91.43% | G08, G09 |
| **Total:** |  | <u>8439</u> | <u>8979</u> | <u>93.99%</u> |  |

**Figure 1:**
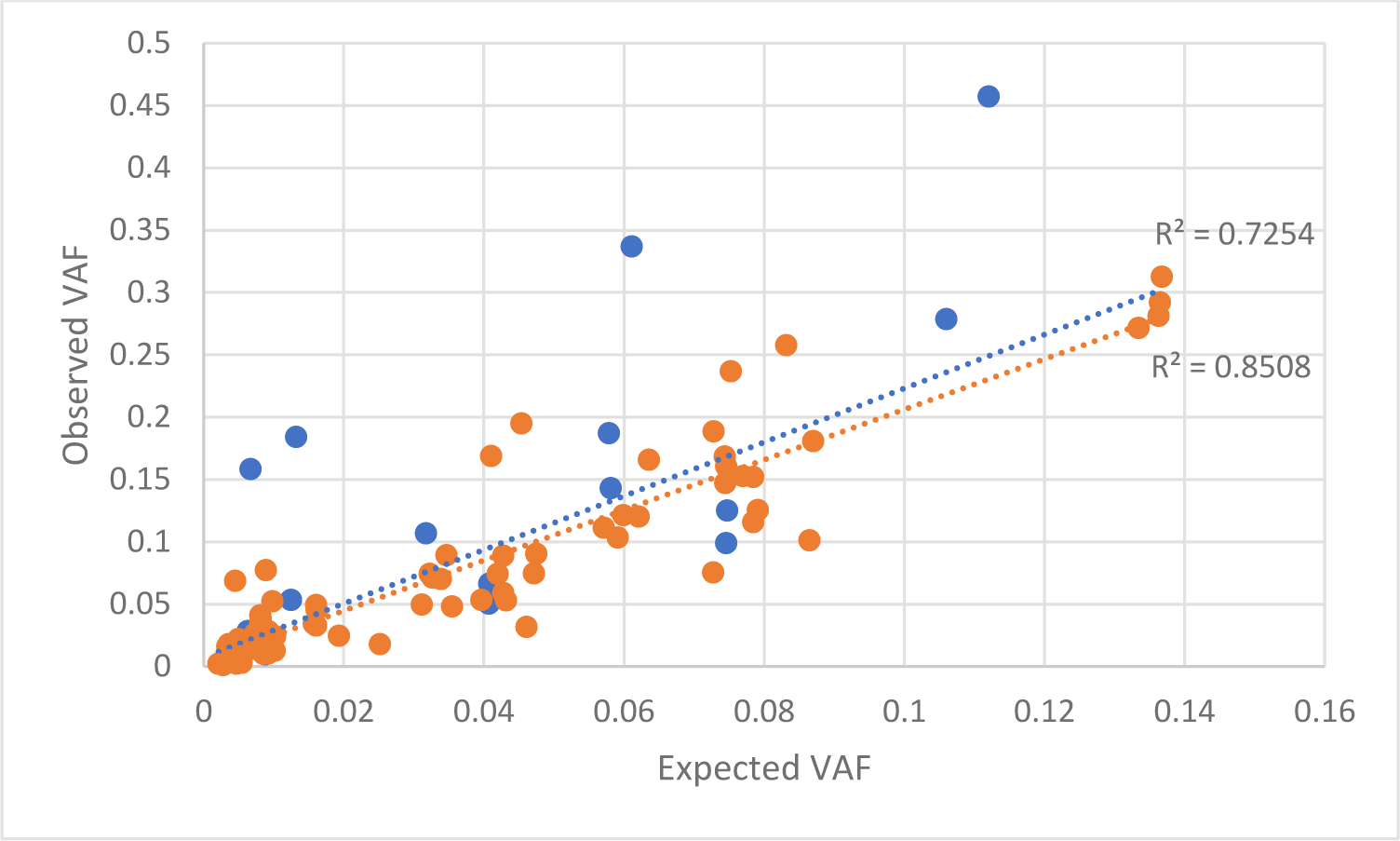
We used serially diluted tumour DNA to obtain expected variant allele frequencies (VAFs) of 0.2-13% for 25 different mutations, including 9 deletions (range 6 – 54 bp in length), one insertion (6 bp), 14 single-base substitutions and one four-base substitution variant. All expected mutations were detected, with a strong correlation between expected and observed VAF for each mutation series (each R^2^ > 0.95), but lower in aggregate over the whole set (R^2^=0.72, blue line). One potential reason for this is preferential amplification of long deletions (viz. short amplicons) compared to the reference allele, and the correlation coefficient does increase (R2=0.85, orange line) when deletions more than 10 bp are not considered. Since preferential detection of long deletions is not an adverse result, we did not further optimise the assay for overall linearity of detection, but avoided quantitative comparison of VAFs across mutations.

### Patient cohort

Patients with histologically confirmed GIST were approached for consent for in a institutional GIST observational study. Serial research blood samples were obtained opportunistically together with routine blood tests The study was approved by the Singhealth Institutional Review Board (Singhealth IRB 2013-107-B). For analyses, patients were grouped into three cohorts, according to the intent of treatment, namely, adjuvant, neoadjuvant and metastatic cohorts.

## Results

### Cohort characteristics

Between 2013 to 2016, 238 blood samples were collected from 65 GIST patients. Patients were analysed according to their disease status as of January 2016. Demographic details are described in Table 2. Of 65 patients, 34, 27 and 4 patients were classified under the metastatic, adjuvant and neoadjuvant cohorts respectively, based upon their treatment intent at point of study entry.

**Table 2:** Cohort demographics

| Demographics |  |
| --- | --- |
| Age (years) |  |
| Median | 65 |
| Range | 25-85 |
| Gender *n* |  |
| Male | 36 |
| Female | 29 |
| Treatment intent |  |
| Adjuvant *n* (%) | 27 (42) |
| Surgery | 27 |
| Imatinib | 18 |
| Neoadjuvant *n* (%) | 4 (6) |
| Imatinib | 4 |
| Surgery | 3 |
| Palliative *n* (%) | 34 (52) |
| Imatinib | 22 |
| Sunitinib | 5 |
| Regorafenib | 3 |
| Primary Mutations *n* |  |
| *KIT* mutations |  |
| Exon 11 | 53 |
| Exon 9 | 4 |
| Exon 13 | 1 |
| Exon 17 | 3 |
| *PDGFRA* mutations |  |
| Exon 18 | 4 |

### Concordance of primary mutations detected in plasma and tissue

Overall, there was 92% concordance of mutations detected in the tissue and plasma (24 out of 26 samples). In two samples, however, the primary mutation detected in plasma was different from that indicated by clinical sequencing. A potential sample mismatch could not be ruled out hence we have removed these timepoints from further consideration. Detection of exon 11 mutations specifically had 89% concordance in tumor tissue and plasma. There was 100% concordance between plasma and tissue samples for exons 9, 13 and 17.

### Metastatic cohort

136 plasma samples from 34 patients with metastatic disease were available for analysis. These patients were grouped according to their radiological disease status. Patients with stable disease (SD) were defined as those who had non-progressive, low volume radiological disease or no radiological evidence of disease throughout all time points of plasma sample collection. Patients with progressive disease (PD) were defined as those who had radiological evidence of disease progression in at least 1 time point of plasma collection. One patient had a single time point plasma sample showing ctDNA at a VAF of 40.5% and this was suspected to be a germline mutation. This patient was removed from further analysis. In this analysis, 17 patients had SD and 16 had PD.

Overall, our assay had a sensitivity of 75% (at least one plasma sample detected in the appropriate setting in 12 out of 16 patients) and a specificity of 94% (16 out of 17 patients) in detecting patient-matched primary mutations in the plasma of patients with metastatic disease. Single or multiple secondary mutations were detected in 13 plasma samples, of which only one sample was taken in a patient who had stable disease to date. The remaining plasma samples were collected either during periods of disease progression or within 6 months prior to disease progression, consistent with the clinical history.

### CtDNA detection in metastatic patients with SD

Among the 17 patients, 16 did not have ctDNA detected at any time point. In low volume, non-progressive disease, we expect minimal ctDNA in the peripheral blood consistent with our finding. One patient had ctDNA detected in one of the 10 timepoints at a very low VAF of 0.05%.

### CtDNA detection in metastatic patients with PD

Of 16 patients defined as having metastatic disease with PD, 12 had at least one plasma sample in which a ctDNA mutation concordant to their primary GIST mutation was detected (Table 3). We demonstrated that ctDNA can be used to (a) detect PD (illustrate that rise in ctDNA levels correspond to radiological disease progression), (b) to monitor treatment response (demonstrate that decline in ctDNA levels correspond to radiological disease response) and (c) to predict PD (show that ctDNA detection predates radiological disease progression) These were illustrated in 7, 4 and 1 out of 12 patients respectively. Secondary mutations were also detected in the appropriate clinical settings.

**Table 3.**
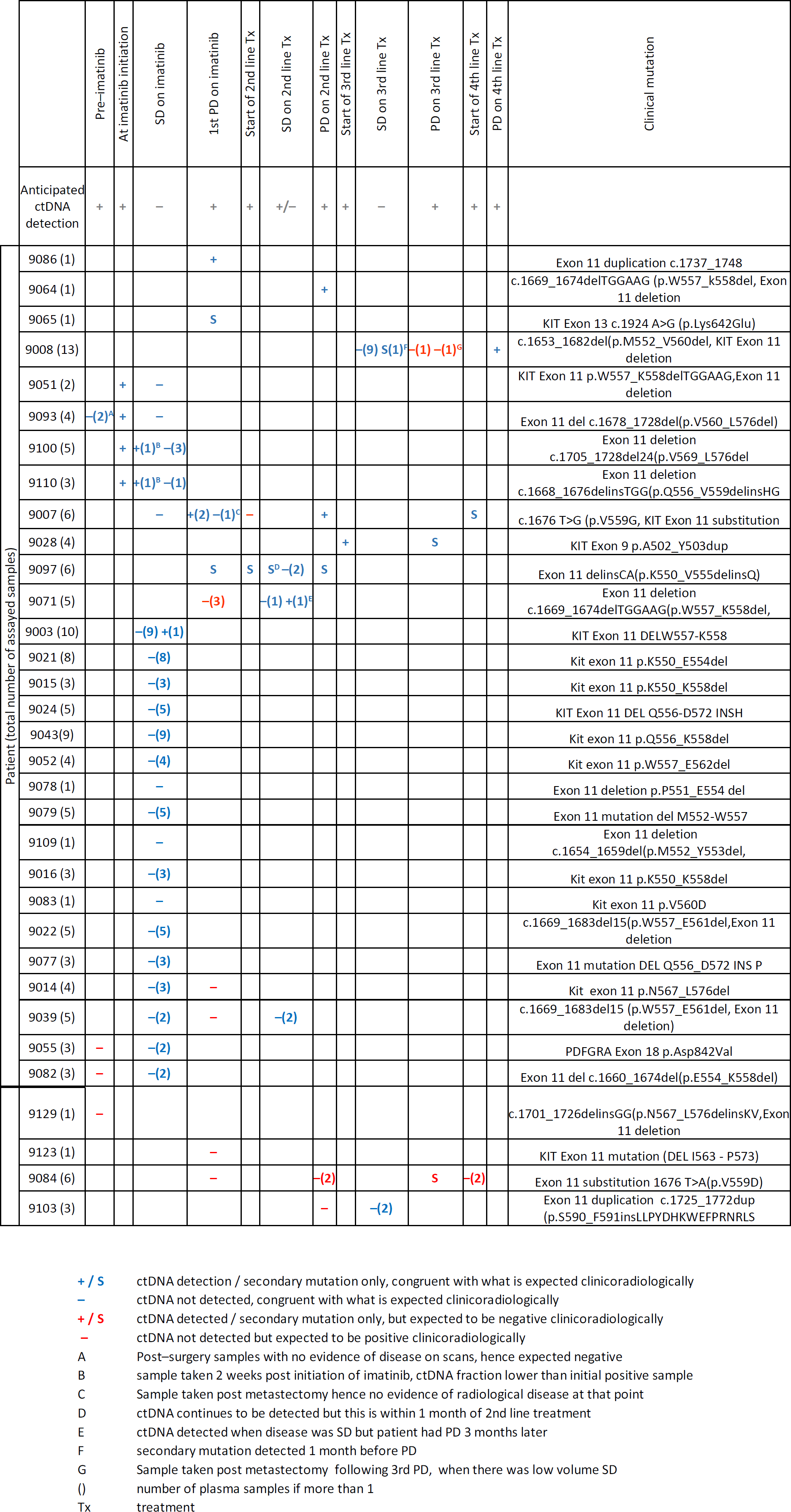

### (a) Detection of PD (Fig 2)

Patient 9007, with *KIT* exon 11 mutated GIST, was on palliative imatinib since 2008. Isolated PD was noted in April 2014, following which metastectomy of the liver lesion was done. ctDNA was detected prior to PD but following surgery, levels remained undetectable till January 2015, when further PD was noted. Imatinib dose was escalated to a maximum of 800mg per day and upon PD, patient was switched to sunitinib and regorafenib, with poor disease response. The ctDNA levels correspondingly demonstrated a rising trend in line with clinical PD. Post PD on imatinib, a secondary mutation in *KIT* exon 17 was detected as well.

**Figure 2:**
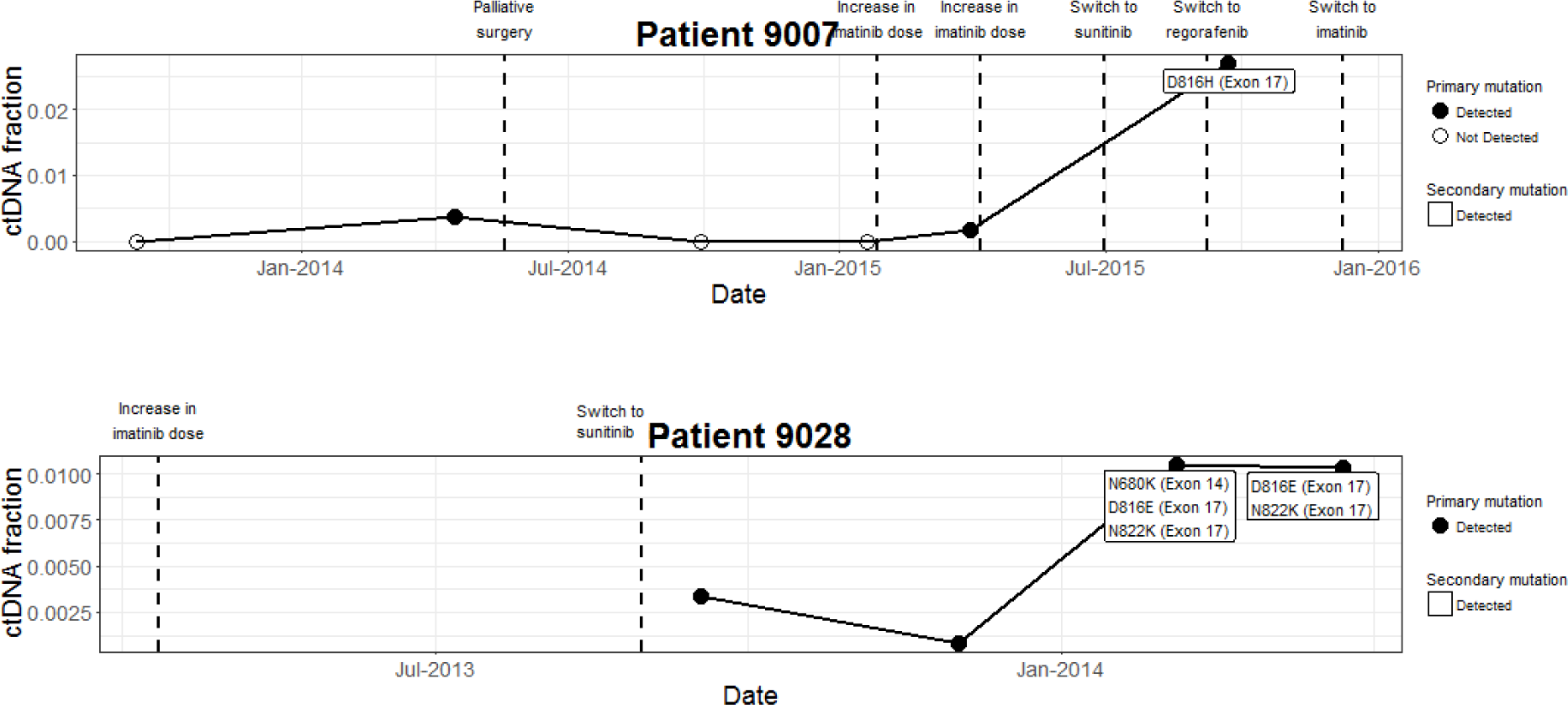
Rise in ctDNA correlates with disease progression.

Patient 9028 was on palliative imatinib since August 2012. PD was noted since April 2013, following which switch to sunitinib was made in August 2013 with a short lived initial radiological response for four months before further PD occurred. Sunitinib was continued beyond progression. Figure 2 illustrated this clinical trajectory of events, with the initial decline of ctDNA amount corresponding to the response to sunitinib, followed by a rising trend in the ctDNA levels. In addition, multiple secondary mutations were also detected at time of progression.

### (b) Monitor Response (Fig 3)

Patient 9100 was started on palliative imatinib in February 2015, prior to which. ctDNA levels were elevated. Within 2 weeks of initiating imatinib, ctDNA levels declined. By 3 months of treatment, ctDNA levels were undetectable.

**Figure 3:**
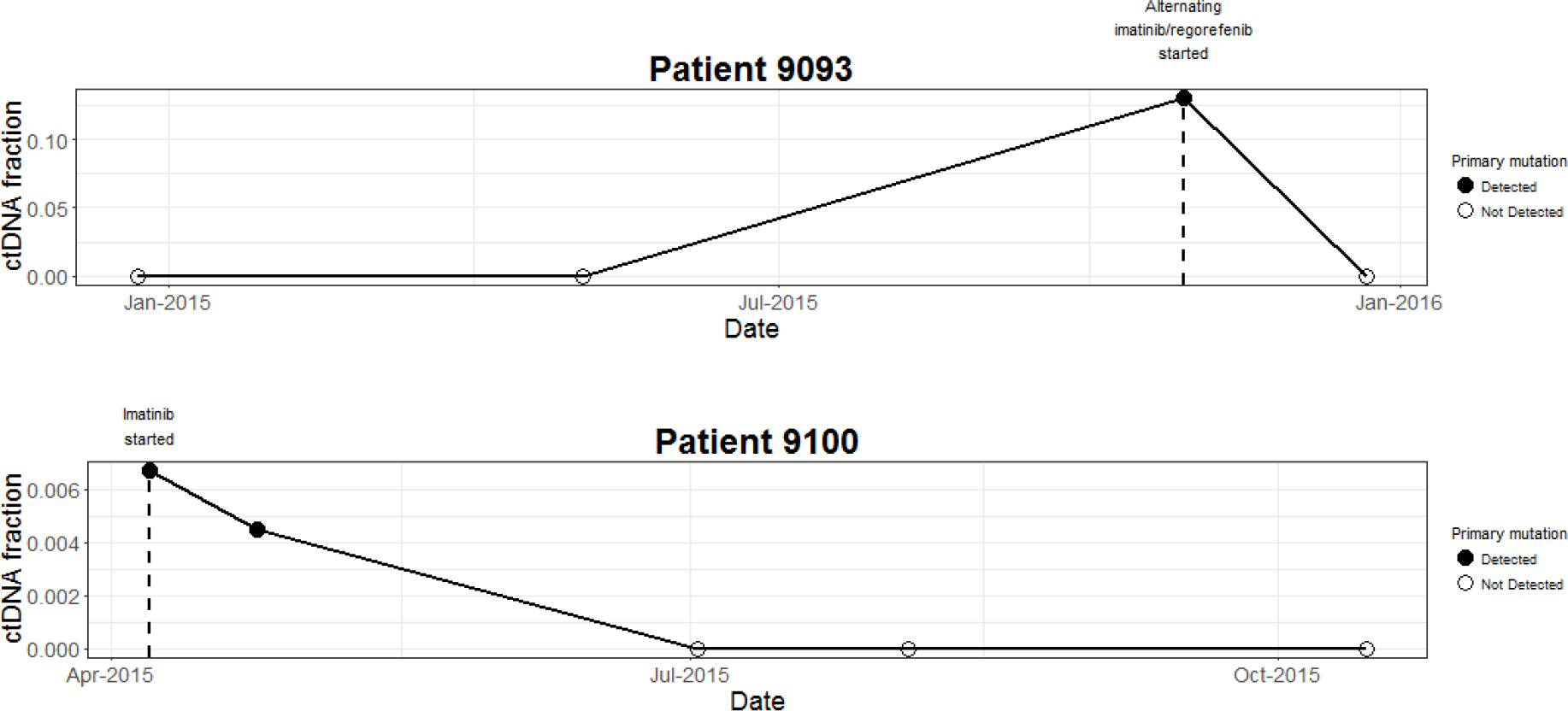
ctDNA as a marker of disease response.

Patient 9093 had metastatic GIST at diagnosis and underwent laparotomy and debulking of tumor in December 2014. There were no plasma samples obtained prior to surgery. Two weeks after surgery, there was no radiological evidence of disease and the plasma samples did not reveal the presence of circulating tumor DNA. The patient became symptomatic with disease recurrence in October 2015 and correspondingly, ctDNA became detectable. Upon initiation of of an alternating imatinib and regorafenib regimen as per clinical trial, ctDNA levels declined, consistent with radiological response noted in this patient.

### (c) Predict PD

Patient 9086 was on imatinib for treatment of advanced GIST but was not compliant to treatment. She underwent palliative surgery for PD in June 2014 following which there was no radiological evidence of disease. A single time point of plasma collection demonstrated detection of ctDNA, and imaging 4 months later demonstrated radiological PD.

In 4 patients (9014, 9084, 9103, 9123) with PD, we could not detect the primary mutation in the plasma samples. In 2 patients (9014, 9084) primary mutation was not detected but one or more secondary mutations were detected at timepoints corresponding to PD. Patient 9103 and 9123 had one plasma sample each taken upon PD to sunitinib and imatinib respectively but ctDNA was not detected. This was postulated to be due to a low amount of ctDNA available for analysis.

### CtDNA was not detected in the adjuvant cohort

Eighty-seven plasma samples were collected from 27 patients with localised GIST at various time points after surgery. None of these patients had ctDNA detected. This was consistent with the absence of active clinical or radiological disease in most of the patients. One patient had disease progression but the last ctDNA sample that was available was 1 year prior to radiological progression.

### CtDNA was not detected in the neoadjuvant cohort

Thirteen plasma samples from 4 patients on neoadjuvant therapy were available. These samples were obtained pre imatinib initiation, pre surgery and post surgery. CtDNA was not detected in any of the samples.

## Discussion

Liquid biopsies have been a subject of great interest in the diagnostic arena for many tumor types. In GISTs, Maier et al. reported using allele specific ligation polymerase chain reaction (PCR) and demonstrated the ability to detect a selected number of pre-specified alterations in *KIT* and *PDGFRa* mutations from plasma^7^. The BEAMing platform is robust in detecting single base substitutions but is less ideal for insertion-deletions, especially in the exon 11 mutations^8^. A recent study using single digital droplet PCR assay demonstrated a high sensitivity of 77% in detecting *KIT* exon 11 mutations. This is a singleplex assay with two drop-off probes that covers 80% of the known exon 11 mutations^9^. These challenges are reflections of the complexities involved in the development of a technology to detect ctDNA, unique to each tumor type. Other studies using NGS based technologies have demonstrated ability to detect ctDNA in the metastatic setting in a small number of patients^10-12^

Our assay has significant advantages as compared to the pre-existing technologies available. The assay was designed to cover a comprehensive spectrum of common mutations seen in GIST, in addition to being able to detect primary and secondary mutations simultaneously. Nearly all the previously reported variants in the exons we target fall within our assay-set, including 99.9% of known exon 11 variants. This is a significant advantage of our assay over some of the pre-existing platforms as the long insertion/deletion mutations in exon 11 have thus far proven to be challenging to detect. In addition, our assay demonstrated 92% concordance of primary mutations detected in tissue and plasma samples. The exon 11 mutations were detected accurately in 89% of patients and there was 100% detection of the rest of the mutations.

Based on the metastatic patients in our cohort, we report a sensitivity of 75% in detecting primary mutations in plasma of metastatic patients. This is higher as compared to other NGS-based technologies, with Hao Xu et al reporting a detection rate of 56.3%^13^ and Kang et al reported a detection rate of 52% using the NGS based technologies^10,11^. This could possibly be attributed to a higher mean sequencing coverage for our assay, reaching 10000 reads per sample in our assay, as opposed to others with a mean coverage of 3000. Also, while our study did not focus on clinical factors that may influence ctDNA levels in plasma, other studies have demonstrated that size of tumor and mutation rate can affect detection of ctDNA in metastatic patients.

In the neoadjuvant cohort, none of the samples had ctDNA detected, including the plasma samples taken prior to initiation of imatinib. Kang et al, however, did demonstrate that in patients with localised GIST, primary mutations were detected in the plasma samples taken prior to surgery, using an NGS based assay^11^. The lack of ctDNA detection in localised disease was echoed in the study by Boonstra et al^9^. Similar to their hypothesis, we postulated that localised GISTs may not shed tumor DNA into the stream as evidenced by the ability of these tumors to grow to large masses locally without having distant metastases.

One of the main limitations of our study is the lack of pre-specified time points for prospective plasma collection as well as defined intervals and paired radiological imaging which would have allowed blood for a more accurate correlation of ctDNA with the clinical course. Additionally, in future studies, standardising the amount of plasma collection would also negate the biases due to sampling errors.

Liquid biopsies can potentially have multiple utilities in the management of GIST patients. In the metastatic setting, a ctDNA assay that has high specificity allows a clinician to minimise frequent radiological imaging when ctDNA remains negative. This might be useful given the prolonged disease free survival in many patients with GIST and could reduce the cumulative number of CT scans required during a patient’s disease course. The ability of ctDNA to predict or demonstrate disease progression accurately either by a rise in the levels of primary mutation or by the development of secondary mutations allows better understanding of development of resistance at a genomic level. With this information, one can then design clinical trials to determine if early intervention at the first emergence ctDNA detectable resistance could improve overall outcome of this disease. A better understanding of resistance mechanisms allows one to better select subsequent therapeutics and explore new strategies to overcome resistance mechanisms including selecting subsequent line treatment based on the drug sensitivities associated with the specific drug resistance mutations that emerge. In the adjuvant setting, the ability to detect minimal residual disease post surgery, may change the paradigm of prognostication for adjuvant treatment. This may then allow for better selection of patients who truly need adjuvant treatment or could allow initiation of imatinib at point of molecular rather than radiologic recurrence.

In conclusion using a novel NGS-based assay that we have developed, we demonstrate the ability to detect primary and secondary circulating *KIT* and *PDGFRA* mutations in GIST patients and showed interesting clinical correlation with the patient’s disease course. Circulating tumor DNA is a powerful tool with potentially diagnostic, predictive and prognostic implications in GIST and warrants validation in a prospective clinical setting.

## References

1. Hirota S, Isozaki K, Moriyama Y, et al: Gain-of-function mutations of c-kit in human gastrointestinal stromal tumors. Science 279:577–80, 1998

2. Heinrich MC, Corless CL, Duensing A, et al: PDGFRA activating mutations in gastrointestinal stromal tumors. Science 299:708–10, 2003

3. Hirota S, Ohashi A, Nishida T, et al: Gain-of-function mutations of platelet-derived growth factor receptor alpha gene in gastrointestinal stromal tumors. Gastroenterology 125:660–7, 2003

4. Heinrich MC, Corless CL, Blanke CD, et al: Molecular correlates of imatinib resistance in gastrointestinal stromal tumors. J Clin Oncol 24:4764–74, 2006

5. Heinrich MC, Maki RG, Corless CL, et al: Primary and secondary kinase genotypes correlate with the biological and clinical activity of sunitinib in imatinib-resistant gastrointestinal stromal tumor. J Clin Oncol 26:5352–9, 2008

6. Liegl B, Kepten I, Le C, et al: Heterogeneity of kinase inhibitor resistance mechanisms in GIST. J Pathol 216:64–74, 2008

7. Maier J, Lange T, Kerle I, et al: Detection of mutant free circulating tumor DNA in the plasma of patients with gastrointestinal stromal tumor harboring activating mutations of CKIT or PDGFRA. Clin Cancer Res 19:4854–67, 2013

8. Demetri GD, jeffers M., Reichardt P., Kang Y.K., Blay J.Y., Rutkowski P.: Mutational analysis of plasma DNA from patients in the phase III GRID study of regorafenib vs placebo in tyrosine kinase inhibitor refractory GIST: correlating genotype with clinical outcomes Journal of Clinical Oncology 31, 2013

9. Boonstra PA, Ter Elst A, Tibbesma M, et al: A single digital droplet PCR assay to detect multiple KIT exon 11 mutations in tumor and plasma from patients with gastrointestinal stromal tumors. Oncotarget 9:13870–13883, 2018

10. Kang G, Bae BN, Sohn BS, et al: Detection of KIT and PDGFRA mutations in the plasma of patients with gastrointestinal stromal tumor. Target Oncol 10:597–601, 2015

11. Kang G, Sohn BS, Pyo JS, et al: Detecting Primary KIT Mutations in Presurgical Plasma of Patients with Gastrointestinal Stromal Tumor. Mol Diagn Ther 20:347–51, 2016

12. Wada N, Kurokawa Y, Takahashi T, et al: Detecting Secondary C-KIT Mutations in the Peripheral Blood of Patients with Imatinib-Resistant Gastrointestinal Stromal Tumor. Oncology 90:112–7, 2016

13. Xu H, Chen L, Shao Y, et al: Clinical Application of Circulating Tumor DNA in the Genetic Analysis of Patients with Advanced GIST. Mol Cancer Ther 17:290–296, 2018

14. Ng SB, Chua C, Ng M, et al: Individualised multiplexed circulating tumour DNA assays for monitoring of tumour presence in patients after colorectal cancer surgery. Sci Rep 7:40737, 2017

15. Somasundaram N LR, Chng J, Ho DL, Gan A, Chua C, Tan Iain BH: An NGS assay strategy with FFPE and cfDNA to determine primary and secondary mutations across the initial diagnosis and subsequent recurrence/progression of patients with localised, recurrent and metastatic GIST. Journal of Clinical Oncology 33, 2015

